# Synergistic and antagonistic effects of chemical pollutants and parasitic fungi on cyanobacterial metabolism

**DOI:** 10.1101/2025.10.07.680941

**Authors:** Erika Berenice Martínez-Ruiz, Jutta Fastner, Stephanie Spahr, Justyna Wolinska

## Abstract

Freshwater communities are increasingly impacted by anthropogenic pollutants, such as the widely used herbicide metolachlor (MET) and chemicals leached from cigarette butts (CBs), one of the most common types of litter worldwide. Cyanobacteria often interact with their parasitic chytrid fungi, which play a role in controlling their growth. Parasites represent an additional biotic stressor that can alter the effects of pollutants on their hosts. However, the metabolic responses of cyanobacteria to simultaneous exposure to pollutants and parasites remain poorly understood. In this study, we investigated the combined effects of abiotic (MET and CB leachate) and biotic (chytrid parasite *Rhizophydium megarrhizum*) stressors on the metabolic response of the toxigenic cyanobacterium *Planktothrix agardhii*. Co-exposure to MET and chytrids led to a synergistic effect, inducing oxidative stress, countered primarily by non-enzymatic mechanisms, whereas MET alone had no measurable effect. In contrast, CB leachate alone induced oxidative stress, but this effect was mitigated when cyanobacteria were also infected by chytrids, indicating an antagonistic interaction. This study demonstrates the complexity of cyanobacterial responses to interacting biotic and abiotic stressors and highlights the importance of considering host-parasite interactions in ecotoxicological assessments of cyanobacteria. A deeper understanding of how pollutants such as MET and CB leachate affect cyanobacteria-chytrid interactions at the metabolic level is crucial for evaluating their broader ecological impacts.

## 1. Introduction

Cyanobacteria are essential primary producers in aquatic system, forming the base of most trophic webs. Eutrophication is a major driver of cyanobacterial proliferation in freshwater systems, often leading to harmful blooms [1]. Climate change can further exacerbate eutrophication and create favorable conditions for cyanobacteria by prolonging water column stratification [2]. These blooms can deplete oxygen levels [3] and produce toxic metabolites that pose significant risks to humans and other organisms [4,5].

Cyanobacterial growth in aquatic environments is regulated by both bottom-up factors, such as nutrient availability [1], and top-down interactions, including parasitism by chytrids [6]. Chytrids, zoosporic fungi from the phylum Chytridiomycota, are common parasites of phytoplankton, including cyanobacteria [7]. Infections by chytrids are lethal, and can delay or suppress bloom formation [6,8,9]. Additionally, chytrids play a key role in aquatic trophic webs by providing alternative links between primary producers and grazers [10–12]. Thus, both chytrids and cyanobacteria are crucial for many ecological and evolutionary processes in aquatic environments.

However, cyanobacteria and chytrids, as well as other aquatic organisms, are increasingly threatened by water pollution, e.g., from agricultural runoff and insufficiently treated domestic and industrial wastewater [13,14]. Common pollutants in aquatic environments include chemicals such as metals, pharmaceuticals, and herbicides like metolachlor (MET), as well as solid waste such as plastics and cigarette butts (CBs), which release a variety of chemicals into the water.

MET, a widely used chloroacetanilide herbicide, is applied with an estimated 100,000 tons annually [15]. In surface waters, concentrations typically range from ng L^-1^ to low μg L^-1^, but can reach up to 400 μg L^-1^ [16–21]. MET has been shown to induce oxidative stress in phytoplankton, likely through the production of reactive oxygen species (ROS), which can alter the activity of antioxidant enzymes such as catalase (CAT), superoxide dismutase (SOD) and glutathione reductase (GR). Moreover, MET has also been linked to lipid peroxidation [22,23] and an increase of cyanotoxin levels and expression of toxin-synthesis genes [24,25]. Such negative impacts on phytoplankton metabolism could impair their survival, growth, and development.

CBs are one of the most commonly littered items worldwide, with around 4.5 trillion improperly discarded annually [26–29]. In urban areas, their density ranges from 1 to 38 CB per m^2^ [30–32]. CBs pose environmental concerns due to the release of microplastics (Green et al., 2023) and chemicals such as nicotine [30,34]. Chemicals released from CBs can adversely affect aquatic organisms, inducing oxidative stress and altering the enzymatic activity and gene expression associated with glutathione *S*-transferase (GST), SOD, and CAT [35,36]. Furthermore, these chemicals can decrease protein levels while increasing lipid concentration and lipid peroxidation [36], leading to metabolic disruptions that impair the survival, growth, and development of aquatic organisms.

The combined effects of biotic and abiotic stressors on aquatic organisms, such as parasites and pollutants, have gained attention in recent years [37,38]. Parasites may exacerbate or mitigate pollutant effects through synergistic, antagonistic, additive, or neutral interactions, depending on the specific host-parasite relationship and type of pollutant involved [37]. For instance, MET promoted cyanobacteria growth in the absence of chytrid parasites, but this effect was lost when both stressors were present [39]. In contrast, CB leachate inhibited cyanobacterial growth alone but promoted it when chytrids were also present [40]. Understanding how hosts respond to combined biotic and abiotic stressors is crucial for assessing their impacts at the ecosystem level.

Biomarkers are biochemical indicators that reflect changes in organisms in response to stimuli, such as exposure to pollutants. These markers provide valuable insights into the metabolic effects of pollutants, which can complement simple growth or inhibition assessments and help elucidate mechanisms of toxicity. Despite their relevance, the metabolic responses of cyanobacteria to simultaneous exposure to chytrid parasites and pollutants remain largely unexplored. Therefore, the goal of this study was to investigate how chytrid parasites and anthropogenic pollutants (MET and CB) affect cyanobacterial metabolism. Specifically, we assessed changes in macromolecule content, oxidative stress biomarkers, and microcystin production in both uninfected and infected cyanobacteria.

Based on previous research, we hypothesized that 1) co-exposure to MET and chytrids enhances the overall cyanobacterial biomarker response compared to MET alone due to synergistic stress; and 2) co-exposure to CB leachate and chytrids elicits a weaker response than CB leachate alone, indicating antagonistic effects.

## 2. Materials and methods

### 2.1 Cyanobacteria and chytrids culturing conditions

The host-parasite system used in this study consists of the toxigenic filamentous cyanobacterium *Planktothrix agardhii* strain NIVA-CYA630 (monoclonal, non-axenic) and its obligate chytrid parasite *Rhizophydium megarrhizum* strain Chy-Kol2008 [41]. *R. megarrhizum* infections are lethal to *P. agardhii*.

Cyanobacterial cultures were maintained in Z8 medium [42] at 16 °C under a continuous light intensity of 20 µmol photons m^-2^ s^-1^. Chytrids were cultivated by transferring zoospores to uninfected cyanobacterial cultures every three weeks under the same temperature and light conditions. Both chytrid and cyanobacterial cultures were maintained under identical temperature and light conditions for the duration of the experiment.

### 2.2 Exposure experiment

We assessed the effect of biotic (chytrid parasites) and abiotic (chemicals) stressors on *P. agardhii*. The concentrations of MET and CB leachate, as well as the sampling time point, were chosen based on previous studies to ensure high cyanobacterial biovolume, active chytrid infection, minimal filament mortality, and a high likelihood of detecting effects on the evaluated biomarkers [39,40].

The experiment consisted of 30 experimental units, including 2 infection conditions (infected and uninfected cyanobacteria) × 2 pollutants (100 µg L^-1^ MET and CB leachate corresponding to 1 CB L^-1^) and 1 negative control (no pollutant), with 5 replicates per treatment. The experiment was conducted in Falcon® 50 mL cell culture bottles with a final volume of 30 mL.

To produce chytrid zoospores, an exponentially growing cyanobacterial culture was infected 10 days prior to the experiment. Afterwards, the infected culture was filtered using a sterile 5 μm nylon mesh and 3 μm polycarbonate membrane [43]. The resulting zoospore suspension was microscopically examined to confirm the absence of cyanobacterial filaments. Zoospore density was quantified using a Sedgewick Rafter chamber under an inverted microscope (Nikon Ti Eclypse).

Before the experiment, cyanobacterial cultures were maintained as exponentially growing semi-continuous cultures for two weeks. The optical density (OD_750nm_) was adjusted to 0.05 three times per week and on the day of exposure before adding MET or CB leachate. This optical density corresponds to approximately 10^4^ filaments mL^-1^. Eight cultures were incubated, with a final volume of 130 mL each. Four of these cultures were infected with 750 zoospores mL^-1^ from the purified zoospore suspension. Infected cultures were incubated for six days to allow infection to be established. After 6 days, infected and uninfected cultures were pooled separately and distributed to the experimental units described above. All experimental units were incubated for 5 days under the conditions described in section 2.1.

### 2.3 Sample collection and processing

For chemical analyses, 1 mL of liquid culture was filtered using 0.22 μm regenerated cellulose syringe filters (Chromafil® Xtra RC-20/13, Macherey-Nagel) to remove bacteria and fungi. Samples were collected on days 0 and 5 from each experimental unit and stored at 4 °C until analysis.

To measure biovolume and parasite fitness proxies, 1 mL of culture was collected from each experimental unit on day 5, fixed with acid Lugol and stored at 4 °C. All sample identities were blinded and randomized before analysis.

To determine total and intracellular microcystin (MC) concentrations in uninfected and infected cyanobacterial cultures, 1 mL of culture was collected from each experimental unit on day 5. The samples were frozen-thawed twice and sonicated for 10 min. To measure extracellular MC concentrations, 1 mL of culture was filtered through a 0.7 µm glass fiber filter (GF/F Grade, Whatmann®). Both sonicated extracts and filtrates were further filtered (PTFE, 0.45 μm, Waters) before analysis. All sample identities were blinded and randomized before analysis.

To quantify additional biomarkers, total biomass was collected from the remaining volume in each experimental unit on day 5 by filtering it through a 0.7 µm glass fiber filters (GF/F Grade, Whatmann®). Biomass was recovered under sterile conditions, resuspended in 1 mL of phosphate buffer (PBS, pH 7.4), and transferred to a 1.5 mL centrifuge tube. The biomass was washed three times with PBS by centrifugation at 15,000 × *g* for 5 min at room temperature. After the final wash, cells were resuspended in 1 mL PBS and disrupted by ultrasonication (Hielscher Ultrasonics®). Ultrasonication was performed in three 30-second cycles at 35% amplitude and 0.5 cycles, while samples were kept on ice to prevent overheating. Between cycles, samples remained on ice. This protocol ensures the disruption of more than 90% of cyanobacterial and chytrid cells (data not shown).

One tablet of cOmplete™ Mini EDTA-free protease inhibitor cocktail (Roche®) was added per 10 mL of PBS used for ultrasonicated samples. After disruption, samples were separated into two fractions. One fraction was centrifuged at 9,000 × *g* for 20 min at 4 °C, and the supernatant (cell-free extract) was collected and used for protein content and enzymatic determinations. The second fraction (crude extract) was not centrifuged and was used to determine thiobarbituric acid reactive substances (TBARS) and lipid content. Sample identities were randomized before analysis.

### 2.5 Recorded parameters

#### 2.5.1 Chemical analyses of MET, nicotine and cotinine

S-metolachlor (MET, CAS 87392-12-9, purity 99.1%) was purchased from Santa Cruz Biotechnology, Inc. A stock solution was prepared in sterile Milli-Q water at a final concentration of 3 mg L^-1^ and stored at 4 °C. MET concentrations were measured to confirm the initial concentration in each experimental unit and to verify its stability throughout the experiment.

The CB leachate stock solution was prepared as described in Guttman et al. (2024) with a final concentration equivalent to 100 CB L^-1^. Due to the complex nature of CB leachate, which is a mixture of different chemicals, assessing the concentration and stability of individual compounds is challenging. We chose nicotine, a major organic compound in CBs, and its metabolite, cotinine, as indicators to monitor the stability of the CB leachate throughout the experiment. While these two compounds provide valuable insights, we recognize that they may not fully represent the stability of all components in the complex CB leachate.

MET, nicotine, and cotinine were quantified using liquid chromatography electrospray ionization tandem mass spectrometry (LC-ESI-MS/MS) with an Agilent 1290 Infinity II UHPLC system coupled to an Agilent 6470 triple quadrupole mass spectrometer in positive ionization mode, following the previously described methods [39,40].

#### 2.5.2 Host biovolume and parasite fitness proxies

To standardize biomarker concentrations across treatments, we measured the biovolume of both cyanobacteria and chytrids in all cultures. This total biovolume measurement was used to calculate biomarker concentrations per mm³ of biomass. Chytrid biomass accounted for less than 1% of the total culture biovolume (0.7 % ± 0.2, Table S1). Therefore, all measured macromolecule concentrations and enzymatic activities were attributed to the cyanobacteria. This approach ensured consistent expression of biomarker data across treatments, while acknowledging potential minor contributions from chytrid biomass.

Cyanobacterial biovolume was quantified using an inverted microscope (Nikon Ti Eclipse). Filament volume was estimated by measuring the filament length in 10 fields of a Sedgewick Rafter chamber per individual sample and using the following formula:

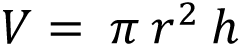

where *r* is the mean radius of cyanobacterial filaments (mean width divided by 2), and *h* is the mean filament length. The mean filament radius was calculated for each exposure treatment (control, MET, and CB leachate) and condition (uninfected and infected) by measuring the width of 10 randomly filaments per sample. Dead cyanobacterial filaments resulting from chytrid infection (i.e., empty, translucent filaments) were excluded from the biovolume calculations.

Total chytrid biovolume was calculated as the sum of sporangia and zoospore volumes, quantified using a Sedgewick Rafter chamber under an inverted microscope (Nikon Ti Eclipse). Sporangia volume was measured in the same fields used for cyanobacterial biovolume estimation. When more than six sporangia were present on the same side of a host cell, their combined volume was estimated as a single ellipsoid, as determining the exact volume of overlapping sporangia poses a challenge. Empty sporangia were excluded from chytrid biovolume calculations. The volume of individual sporangia was calculated using the formula, considering them as single rotational ellipsoids:

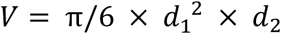

where *d_1_* and *d_2_* are the short and long semi-axes, respectively.

Zoospores volume was determined by quantifying the zoospore density in the same fields used for cyanobacterial biovolume measurements. Based on unpublished data, each zoospore was estimated to have an average volume of 15.5 μm³.

Additionally, we assessed two parasite fitness proxies: infection prevalence and intensity, to determine the level of infection in the cyanobacterial culture and to identify potential effects of pollutants on chytrid infection dynamics. Infection prevalence indicates the proportion of infected filaments within a host population and was calculated as the percentage of cyanobacterial filaments containing encysted zoospores or sporangia after randomly examining 200 filaments under an inverted microscope (Nikon Ti Eclipse). Infection intensity represents the number of parasites infecting a single host. It was determined as the average number of parasites (encysted zoospores or sporangia) per infected filament after randomly examining 50 infected filaments per sample. When more than six infections were present on the same side of a single host cell, infected filaments were categorized as having six infections, as accurately counting zoospores or sporangia above this threshold is challenging due to overlapping structures.

#### 2.5.3 Biomarkers Proteins

Protein content was quantified using the Bradford method [44]. Measurements were made in 96-well transparent flat-bottom plates adding 95 µL of Bradford reagent and 5 µL of cell-free extract per well. Absorbance was measured at 595 nm using bovine serum albumin as a standard. Each sample was quantified six times as technical replicates. Protein content was adjusted to the total biovolume and expressed as mg mm^-3^.

##### Lipids

Lipid content was determined using the sulfo-phospho-vanillin method [45]. Lipids were first extracted by adding 300 µL of a chloroform:methanol solution (2:1, v/v) to 100 µL of crude extract [46]. After mixing by inversion, 250 µL of water were added to induce phase separation. The lower phase was transferred to a 1.5 mL centrifuge tube and heated at 80 °C to evaporate the solvent. Following evaporation, 25 µL of concentrated sulfuric acid were added, and samples were heated for 10 min at 80 °C. Samples were then cooled to room temperature, and 250 µL of phosphor vanillin reagent (1.2 g L^-1^ vanillin in 68% phosphoric acid) were added. After gentle mixing and incubation on ice for 5 min, 100 µL of each sample were transferred to a 96-well flat-bottom transparent microplate, and absorbance was measured at 525 nm. Canola oil was used as standard [47]. Each sample was quantified three times as technical replicates. Lipid content was adjusted to the total biovolume and expressed as mg mm^-3^.

##### Microcystins (MCs)

Microcystins (MCs) concentrations were determined by liquid chromatography–tandem mass spectrometry (LC–MS/MS) using an Agilent 1290 series HPLC system (Agilent Technologies, Waldbronn, Germany) coupled to an API 5500 QTrap mass spectrometer (AB Sciex, Framingham, MA, USA) equipped with a turbo-ion spray interface. Samples were separated at 30 °C using a Purospher STAR RP-18 end-capped column (30 mm x 4 mm, 3 mm particle size, Merck, Germany) with a mobile phase consisting of 0.5% formic acid (A) and acetonitrile with 0.5% formic acid (B) at a flow rate of 0.5 mL min^-1^ with a gradient program (0 min 25% B, 10 min 70% B, 11 min 70% B). The injection volume was 10 μL. Identification and quantification of MCs ([Asp3]-MC-RR, MC-RR, MC-YR, [Asp3]-MC-LR, MC-LR, MC-LW, MC-LF and MC-LA; standards purchased from Novakits, France) was performed in the Multiple Reaction Monitoring (MRM) mode. The detection limits of the different congeners ranged from 0.05 to 0.3 μg L^-1^. Intracellular MC (MCi) concentration was estimated as the difference between total MC (MCt) and extracellular MC (MCe). Among the MCs analyzed, we exclusively detected [Asp3]-MC-RR, [Asp3]-MC-LR, and an additional MC variant present at a concentration too low for precise identification. To maintain clarity throughout the text, these will be referred to as DMC-RR, DMC-LR, and MC-X, respectively.

##### Thiobarbituric acid reactive substances (TBARS)

TBARS, an indicator of lipid peroxidation levels, were quantified according to Ohkawa et al. (1979). First, 400 μL of the reaction solution (0.375% 2-thiobarbituric acid, 0.5% sodium dodecyl sulfate, and 9.375% acetic acid pH 4) were added to 100 μL of crude extract. The mixture was incubated at 95 °C for 1 h. After cooling on ice, samples were centrifuged at 1,200 × *g* for 5 min. Finally, 150 µL of each sample were transferred to a 96-well microplate, and absorbance was measured at 532 nm. 1,1,3,3,-tetramethoxypropane (TMOP) was used as standard. Each sample was quantified four times as technical replicates. Since other molecules besides malondialdehyde and 4-hydroxyhexenal (the primary products of lipid peroxidation) can also react with thiobarbituric acid, the measurements are referred to as TBARS levels. The TBARS content was adjusted to the total biovolume and expressed as μmol mm^-3^.

##### Superoxide dismutase (SOD) activity

SOD activity was quantified by a colorimetric technique using the SOD activity assay kit (Sigma-Aldrich®). Briefly, 20 μL of cell-free lysate and 160 μL of working solution were added to a 96-well plate. The reaction was initiated by adding 20 μL of xanthine oxidase working solution to each well, and the plate was incubated at 20-25 °C for 30 min. Absorbance was measured at 450 nm. SOD supplied by the manufacturer was used as standard. The SOD activity was adjusted to the total biovolume and expressed as units mg^-1^ of protein.

##### Glutathione S-transferase (GST) activity

GST activity was quantified using a modified version of the method described by Nauen and Stumpf (2002). In a black-walled, clear flat-bottom 96-well microplate, 100 μL of monochlorobimane (500 µM), 40 μL of PBS (pH 7.4) and 10 μL of cell-free extract were added per well. The enzymatic reaction was initiated by adding 100 μL of reduced glutathione (GSH, 0.1 mM). Fluorescence intensity was measured at 460 nm (excitation: 355 nm) at the start of the reaction and then every minute for 5 min. GSH was used as standard. GST activity was calculated as the rate of GSH conversion per minute and expressed as μmol min^-1^ g^-1^ of protein.

### 2.6 Data analyses

To integrate biomarker results and evaluate the global response of the host under multiple stressors, we calculated the Integrated Biomarker Response version 2 (IBRv2) following the method described by Sanchez et al. (2013). This analysis compares individual biomarker data to a reference mean, which, in our study, corresponds to the mean biomarker values of non-exposed cultures (for both uninfected and infected conditions, respectively). To preserve individual data variability, we included biomarker values from all five replicates of the control, MET-exposed, and CB leachate-exposed treatments. For the IBRv2 analysis of uninfected cultures, the extracellular MC-X values were excluded due to the majority of samples being below the detection limit, which resulted in numerous zero values that were incompatible with the IBRv2 analysis. For the infected cultures, the extracellular DMC-LR and MC X were excluded for the same reason. IBRv2 values closer to those of the unexposed group indicate a more similar physiological state, while values that deviate further from the control group represent a more divergent response relative to the control condition. Biomarker values are displayed in a star plot, where positive values indicate biomarker increase and negative values indicate biomarker inhibition relative to the control.

Linear models were used to assess the effects of multiple stressors on parasites fitness proxies, biomarker responses, and IBRv2. For the cyanobacterial biovolume, a linear mixed model was used to account for repeated measurements within the same sample (multiple observations). A two-way interaction was included in the IBRv2 model to test for interactions between treatment (control, MET or CB leachate) and condition (uninfected and infected). The significance of this interaction was assessed by comparing a model with the interaction term to one without it.

Residuals distribution was visually examined to ensure normality and homogeneity of variance. If assumptions were not met, data were log- or square root-transformed to normalize residuals. Homoscedasticity of residuals was tested with the Breusch Pagan test. When transformations failed to meet assumptions, non-parametric linear models were applied.

### 2.7 Software for data processing, analysis, and visualization

Biovolume measurements were performed using the NIS-Elements BR 4.5 software (Nikon®). Statistical analyses and figures were carried out in R Statistical language (version 4.2.3; R Core Team, 2023), using the packages described below. Linear models were run using base R, the linear mixed model was run using *lme4* [51], and post-hoc tests were conducted with *emmeans* [52]. Data visualization for normality and homogeneity of variance was done with *ggResidpanel* [53], while the Breusch-Pagan test was performed using *lmtest* [54]. Conditional and marginal *R^2^* for each mixed model was estimated according to Nakagawa and Schielzeth [55] with *MuMIn* [56]. The IBRv2 was calculated using the *IBRtools* package [57]. Figures were made with *ggplot* [58], *ggpubr* [59], *cowplot* [60], and *fmsb* [61]. Additionally, various *tidyverse* packages [62] were used for data import, export, tidying, and organization. R code used for data analysis and generation of results is provided as Supplementary material.

## 3. Results

### 3.1 Chemical analyses of MET, nicotine and cotinine

We measured the MET concentration at the start and end of the experiment to assess its stability in the cultures. The initial MET concentrations were 95.1 and 92.4 μg L^-1^ in the uninfected and infected cultures, respectively (Figure S1A), which was close to the target concentration of 100 μg L^-1^. MET remained stable throughout the experiment, indicating no significant abiotic or biotic transformations under the tested conditions (Table S2). For simplicity, the target concentration is discussed in the following sections.

To assess the stability of CB leachate, we measured nicotine and cotinine concentrations as proxies. The initial nicotine concentrations were 336.1 and 339.4 μg L^-1^ in uninfected and infected cultures, respectively (Figure S1B). The nicotine concentration decreased slightly by 8.7% in uninfected and 5.4% in infected cultures, indicating partial nicotine transformation (Table S2).

The initial cotinine concentrations were similar in both culture types: 5.3 μg L^-1^ in uninfected and 5.4 μg L^-1^ in infected cultures. After five days, cotinine levels increased by 11% in uninfected cultures and 61% in infected cultures (Figure S1C). These changes correspond to cotinine yields, representing the fraction of nicotine converted to cotinine, of 1.9% and 16.8%, respectively. This increase, particularly in infected cultures, suggests partial nicotine transformation to cotinine due to abiotic or biotic factors under the tested conditions (Table S2).

### 3.2 Exposure experiment

In uninfected cultures, the cyanobacterial biovolume increased slightly under MET and CB leachate exposure, but the effect was not significant (Figure S2A). In infected cultures, the biovolume was significantly higher under MET exposure compared to the control (*p* = 0.0354), while CB leachate exposure had no effect. Treatment (MET or CB leachate) accounted for 16% of the total variance in the model (conditional *R^2^* - marginal *R^2^*; Table S3).

Infection prevalence was significantly lower in cultures exposed to CB leachate compared to both control and MET-exposed cultures, indicating that CB leachate hindered chytrid infection (adj. *R^2^* = 0.65; Control - CB: *p* =0.0047; MET - CB: *p* = 0.0009; Figure S2B, Table S4). However, infection intensity was similar in all treatments compared to the control (Figure S2C, Table S4).

Protein and lipid concentrations in cyanobacteria exposed to CB leachate alone were significantly lower than in the control (proteins: adj. *R^2^*= 0.40, *p* = 0.0161; lipids: adj. *R^2^* = 0.47, *p* = 0.0068; Figures 1A and 1B; Table S4), while MET exposure had no effect. In cultures exposed to both chemical stressors (MET or CB leachate) and parasites, protein and lipid levels were comparable across treatments.

**Figure 1.**
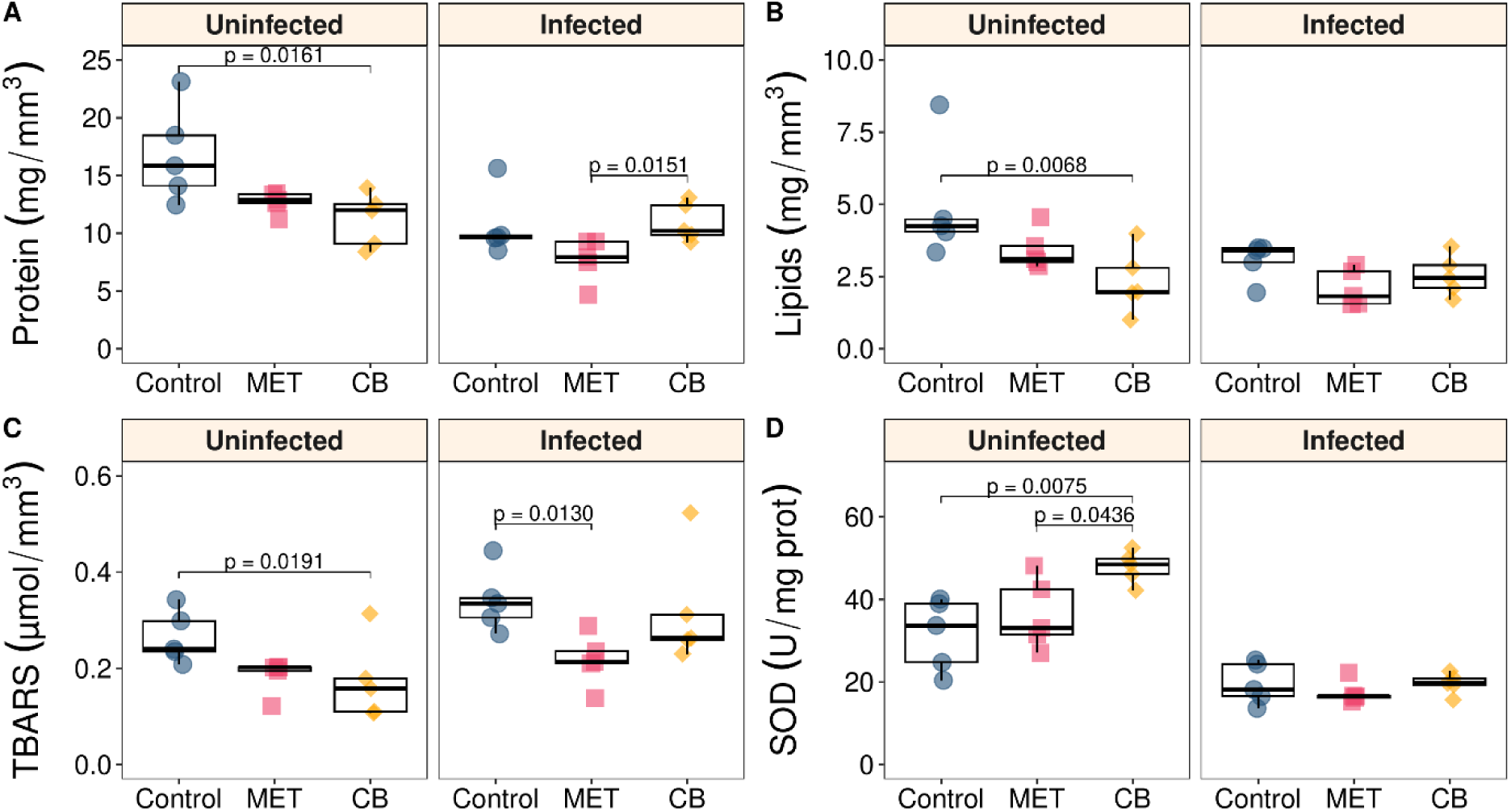
Biochemical parameters and oxidative stress markers in cyanobacteria exposed to chemical pollutants with and without chytrid parasites. A) Protein concentration, B) lipid concentration, C) thiobarbituric reactive substances (TBARS) concentration, and D) superoxide dismutase (SOD) activity. Protein, lipid and TBARS concentrations were adjusted to the total biovolume of each replicate per treatment and condition. SOD activity was adjusted to the protein content of each replicate per treatment and condition. Boxes indicate upper and lower quartiles, the dark middle line indicates the median, whiskers indicate 1.5 times the interquartile range and black points represent outliers (*n* = 5). MET: metolachlor; CB: cigarette butt leachate.

TBARS levels in uninfected cultures were unaffected by MET but significantly decreased under CB leachate compared to the control (adj. *R^2^*= 0.41, *p* = 0.0191; Figure 1C; Table S4). In infected cultures, MET exposure significantly reduced TBARS levels compared to the control (adj. *R^2^* = 0.42, *p* = 0.0130), while CB leachate had no effect (Figure 1C; Table S4).

Three different MCs were detected in *P. agardhii* strain NIVA-CYA630. The dominant variant was DMC-RR (90-95% of total MCs), followed by DMC-LR (4-9%), and an unidentified variant, designated as MC-X (< 1%) (Table S5). Total and intracellular DMC-RR and DMC-LR concentrations followed similar patterns (Figures 2 and S3). In uninfected cultures, MET exposure did not affect their concentrations, whereas CB leachate exposure significantly reduced their levels (DMC-RRt Control-CB: adj. *R^2^* = 0.34, *p* = 0.0304; DMC-LRt Control-CB: adj. *R^2^* = 0.42, *p* = 0.0151; DMC-RRri Control - CB: adj. *R^2^* = 0.34, *p* = 0.0304; DMC-LRi Control-CB: adj. *R^2^* = 0.42, *p* = 0.0151, Table S6). Conversely, in infected cultures, MET exposure significantly decreased total and intracellular DMC-RR and DMC-LR concentrations compared to the control, while CB leachate exposure had no effect (DMC-RRt Control - MET:, adj. *R^2^* = 0.55, *p* = 0.0258; DMC-LRt Control-MET: adj. *R^2^* = 0.50, *p* = 0.0045; DMC-RRi Control - MET: adj. *R^2^* = 0.55, *p* = 0.0258; DMC-LRi Control-MET: adj. *R^2^* = 0.48, *p* = 0.0056, Table S6).

**Figure 2.**
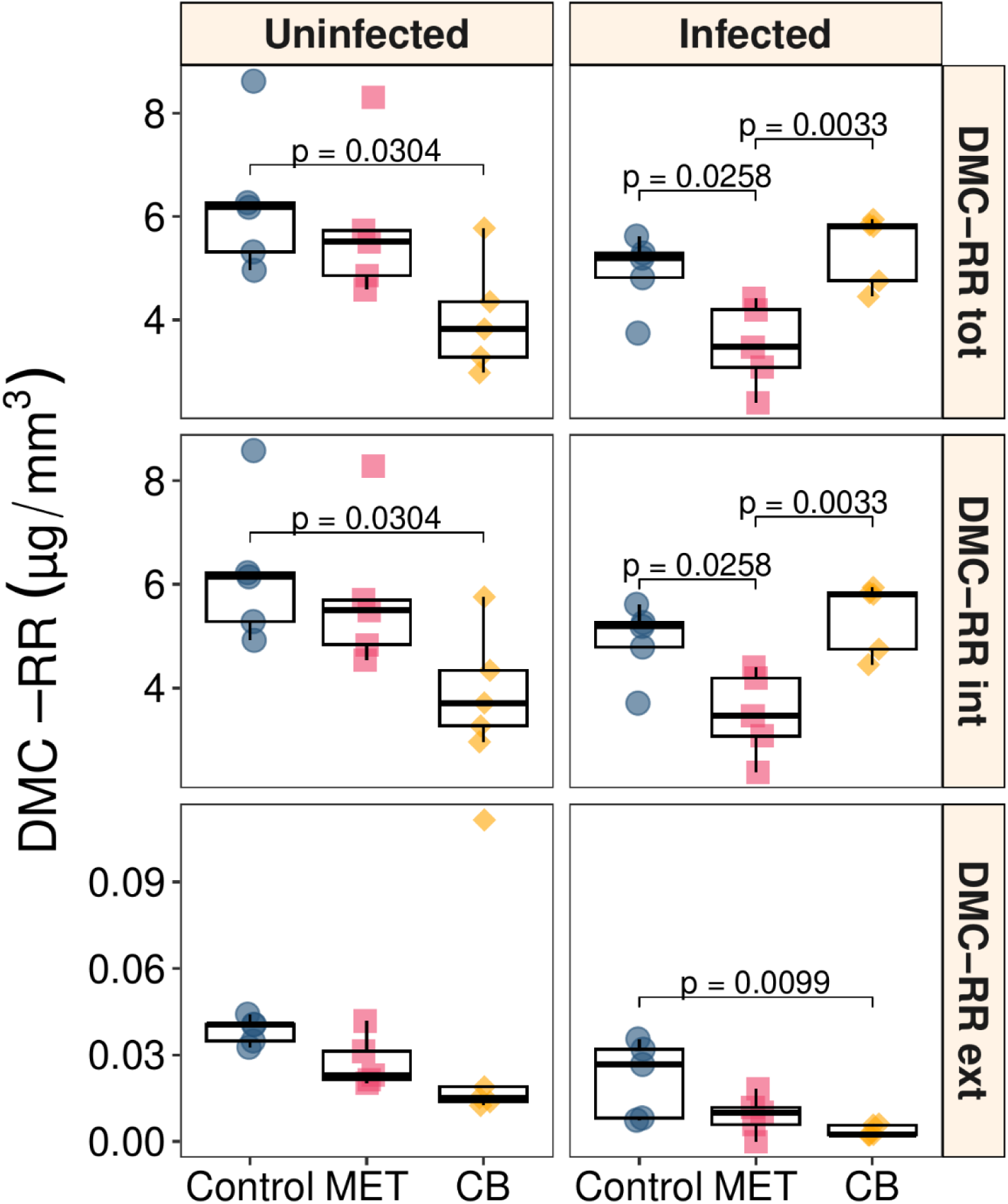
Total, intracellular and extracellular DMC-RR concentrations in cyanobacteria exposed to chemical pollutants with and without chytrid parasites. DMC-RR concentrations were adjusted to the cyanobacterial biovolume of each replicate per treatment and condition. Boxes indicate the upper and lower quartiles, the dark middle line indicates the median, whiskers indicate 1.5 times the interquartile range and black points represent outliers (*n* = 5). MET: metolachlor; CB: cigarette butt leachate.

Extracellular DMC-RR and DMC-LR levels in uninfected cultures were consistent across treatments (Figures 2 and S3). In infected cultures, exposure to CB leachate significantly reduced extracellular DMC-RR levels (DMC-RRe Control-CB: adj. *R^2^* = 0.44, *p* = 0.0099; Table S6) and both CB leachate and MET exposure reduced extracellular DMC-LR concentrations relative to the control (DMC-LRe Control - CB & Control - MET: adj. *R^2^* = 0.41, *p* = 0.0300; Figure 2 and S3; Table S6).

Total and intracellular MC X concentrations were significantly lower in uninfected cultures exposed to CB leachate compared to the control (MC Xt-i Control-CB: adj. *R^2^* = 0.47, *p* = 0.0098; Figure S4; TableS6), but there were no differences in infected cultures. Extracellular MC X concentrations were consistent across treatments regardless of infection condition (Figure S4; TableS6).

SOD activity in uninfected cultures decreased significantly under CB leachate exposure compared to the control (adj. *R^2^* = 0.48, *p* = 0.0075, Figure 1D; Table S4), but was unaffected by MET exposure. Infected cultures had similar SOD activity across all treatments.

GST activity remained consistent across all treatments, regardless of parasite presence or exposure to MET or CB leachate (Figure S5, Table S4).

### 3.3 IBRv2

The star plots of the A index for each evaluated biomarker, showing deviation from the control reference, are presented in Figures 3A and 3B. In uninfected cultures, most biomarkers in MET-exposed cyanobacteria were close to control values, whereas those from CB leachate-exposed cultures showed greater deviations, primary indicating biomarker reductions (Figure 3A). In contrast, in infected cultures, MET exposure resulted in larger biomarker deviations, with most showing a reduced response, while biomarkers in CB leachate-exposed cultures remained close to control values (Figure 3B).

**Figure 3.**
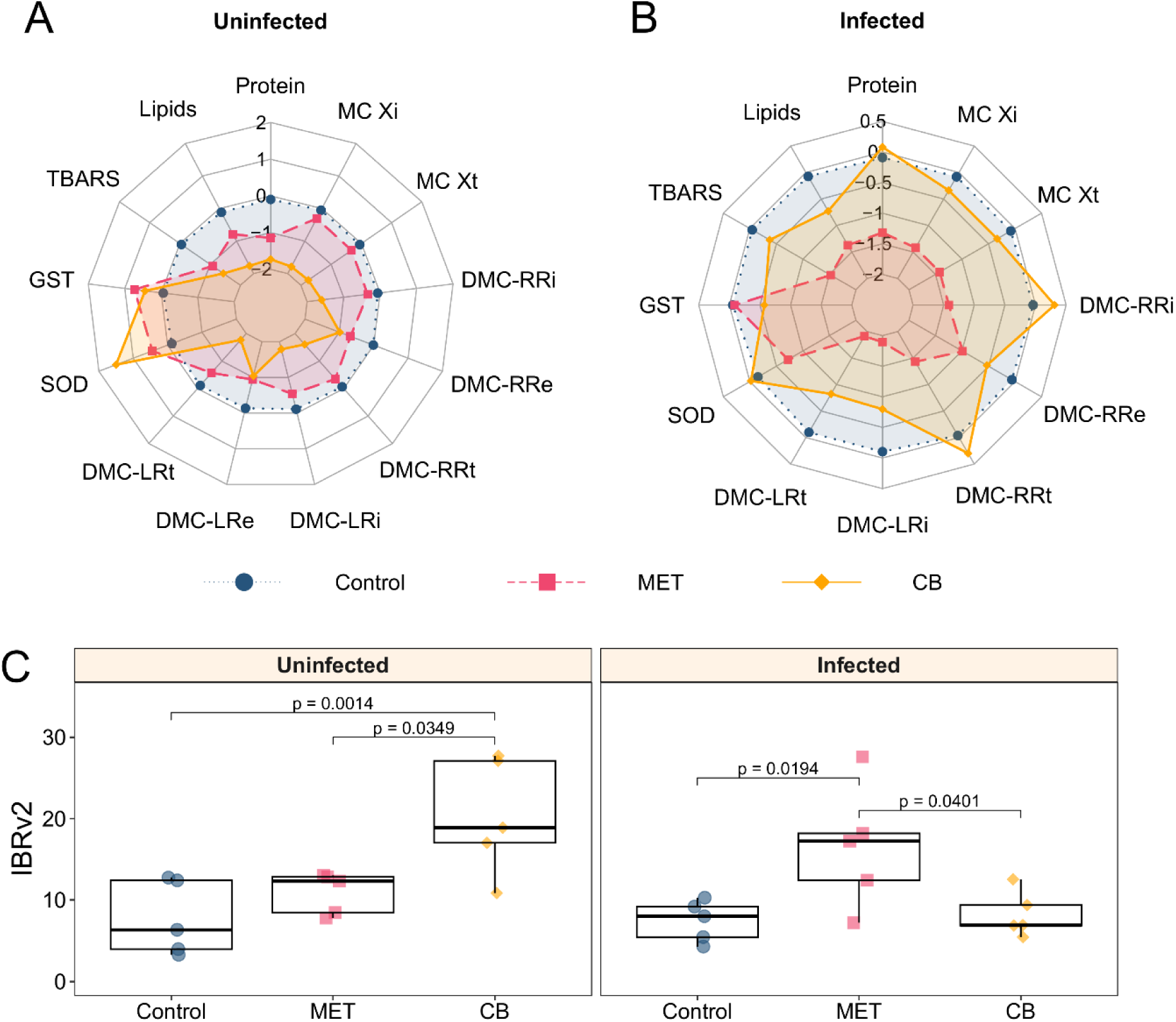
Integrated Biomarker Response version 2 (IBRv2) in cyanobacteria exposed to chemical pollutants with and without chytrid parasites. Star plot of biomarker deviation index of A) uninfected and B) infected cultures; C) comparison of IBRv2 values between treatments. In C, boxes indicate the upper and lower quartiles, the dark middle line indicates the median, whiskers indicate 1.5 times the interquartile range and black points represent outliers (*n* = 5). MET: metolachlor; CB: cigarette butt leachate; MC t: total microcystin; MC i: intracellular microcystin, MC e: extracellular microcystin; SOD: superoxide dismutase; GST: glutathione *S*-transferase; TBARS: thiobarbituric acid reactive substances.

IBRv2 values provide an insight into how closely treated cultures resemble the control response. In uninfected cyanobacterial cultures, IBRv2 values were comparable to the control with MET exposure but significantly higher with CB leachate exposure (adj. *R^2^* = 0.45, *p* = 0.0014; Figure 3C; Table S2). In infected cultures, IBRv2 values were significantly higher than the control when exposed to MET (adj. *R^2^* = 0.45, *p* = 0.0194, Figure 3C; Table S2), but exposure to CB leachate resulted in values similar to the control. The difference in IBRv2 between uninfected and infected cultures exposed to CB leachate was significant (adj. *R^2^* = 0.45, *p* = 0.0012), suggesting distinct overall responses depending on parasite presence.

## 4. Discussion

Our results show that the simultaneous exposure of the cyanobacterium *P. agardhii* to chemical and biological stressors produced distinct metabolic responses, depending on the type of chemical stressor. Exposure to the herbicide MET did not affect any biomarker in uninfected cyanobacteria. However, co-exposure to MET and chytrids led to reduced levels of TBARS and MC in infected cultures, suggesting intensified metabolic stress. Conversely, exposure to CB leachate alone decreased biomolecule concentrations and enzymatic activity in uninfected cyanobacteria but had no effect on infected cultures. This suggests that chytrid parasites mitigate the metabolic stress induced by CB-released chemicals. Overall, our findings highlight how pollution by single pollutants like MET or complex chemical mixtures like CB leachate can produce distinct metabolic responses when combined with biological stressors such as chytrid parasites.

Chemical analysis confirmed the stability of MET, consistent with previous work [39]. In contrast, nicotine concentrations decreased by 9% in uninfected cultures and by 5% in infected cultures, suggesting a slow nicotine transformation. Cotinine concentrations increased in both culture types, with a 5.5 times greater increase in infected cultures likely due to the partial transformation of nicotine to cotinine. Several nicotine transformation pathways have been described in bacteria and fungi, leading to the production of diverse metabolites, including cotinine [63]. Considering that both types of cultures were incubated under identical conditions, the observed differences in cotinine formation might be due to different transformation mechanisms induced by cyanobacteria, chytrids, or associated bacteria.

Cyanobacterial biovolume remained unchanged in uninfected cultures exposed to an environmentally relevant concentration of MET (100 µg L^-1^) but increased in infected cultures. This contrasts with our previous findings, where continuous MET exposure promoted cyanobacterial growth, but this effect was reversed by chytrid infection [39]. These differences may be due to the experimental design, as the current study used a single sampling point, whereas the previous study analysed multiple time points, which may allow a better assessment of cyanobacterial growth. Meanwhile, CB leachate (1 CB L^-1^) had no effect on cyanobacterial biovolume in either culture type, which contrasts with other studies showing growth inhibition of aquatic photosynthetic microorganisms by CB leachate [35,40,64]. The discrepancy may be related to differences in exposure duration or concentration.

Neither MET nor CB leachate decreased infection intensity. However, CB leachate reduced infection prevalence, supporting previous findings from exposure of chytrid-infected cyanobacterial cultures to polystyrene nanoplastics [65] and a lower concentration of CB leachate [40]. The latter study hypothesized that under CB leachate exposure, chytrids remain metabolically capable of infecting the host, but their ability to efficiently complete the infection cycle is compromised, resulting in lower infection prevalence without affecting infection intensity [40].

Protein content remained unaffected by MET in both culture types, contrasting with its known mechanism of inhibiting protein synthesis in plants [66]. The effects of MET on protein levels in non-target organisms appear inconsistent, with some studies reporting decreases, such as in the aquatic midge *Chironomus tentans* [67], and others reporting increases, such as in two aquatic plants [68]. In contrast, CB leachate reduced protein levels only in uninfected cyanobacteria, with no effect in co-exposed cultures. Likewise, chemicals released from CBs have been shown to decrease plasma protein content in Nile tilapia [36].

Given that proteins are essential for metabolism, structural integrity, and cell-to-cell communication in cyanobacteria, their reduction may compromise survival, growth, and adaptability to environmental conditions. The exact mechanism remains unclear but may involve the inhibition of protein synthesis, disruption of protein turnover, use of existing proteins as an emergency nitrogen source, or interference with nutrient uptake. These factors could ultimately disrupt the general metabolism of cyanobacteria, as suggested for other pollutants with a similar effect on protein content [36,69,70].

Lipid levels remained unchanged in both uninfected and infected cultures exposed to MET, despite the herbicide’s known role in inhibiting fatty acid synthesis [71]. One explanation for this discrepancy could be that MET alter the concentration of specific lipids synthesized by cyanobacteria rather than reducing total lipid content, as observed in other aquatic photosynthetic organisms [72,73]. For instance, MET alters the concentrations of palmitic acid (16:0) and linoleic acid (18:2(n-6)) in the freshwater diatom *Gomphonema gracile* [73]. Interestingly, these two fatty acids are also present in *P. agardhii* and may play a role in chytrid metabolism, as they have been detected in chytrid zoospores following infection of *P. agardhii* [12]. In this context, changes in lipid composition may not affect the growth of uninfected cyanobacteria. However, in infected cyanobacteria, such changes could indirectly affect nutrient availability for chytrids, potentially reducing their fitness over time, as previously reported [39]. Further studies are needed to determine MET’s impact on lipid composition in cyanobacteria.

Similar to its effect on proteins, CB leachate reduced lipid content in uninfected cyanobacteria but had no impact on infected ones. Lipids play essential roles in the metabolism, membrane structure, and energy storage. Under various stress conditions, including exposure to pollutants, aquatic organisms, such as microalgae, cyanobacteria and fish often increase lipid synthesis and energy storage [46,69,74,75]. However, uninfected cyanobacteria in this study showed reduced lipid content, suggesting that CB leachate disrupts lipid biosynthesis or energy storage pathways.

Total and intracellular MC and TBARS concentrations were unaffected by MET in uninfected cyanobacteria. This differs from previous studies showing that pollutants, including MET, 2,4-dichlorophenoxyacetic acid, and nickel (Ni^2+^), increase MC levels in the cyanobacterium *Microcystis aeruginosa* [24,25,46,76]. However, co-exposure to MET and chytrids decreased MC levels. Notably, MCs are known to scavenge reactive oxygen species (ROS) and can be reduced when high ROS levels are present in the cell [77,78]. Moreover, cyanobacteria exposed to MET and chytrids showed a reduced concentration of TBARS, suggesting a decreased in lipid peroxidation in the cells. Chytrid infections may destabilize cyanobacterial membrane and cell walls, which serve as the first physical barriers to infection. Considering that MET may alter lipid composition by reducing PUFA levels (the primary lipid type affected by lipid peroxidation [79]), we hypothesize that the lower TBARS concentrations could be attributed to two factors: a possible reduction in PUFA synthesis due to MET exposure, and a reduced membrane surface area available for oxidation due to chytrid infection. The reduction in MCs and TBARS concentrations, alongside unchanged SOD and GST activity, suggests that non-enzymatic antioxidant mechanisms counteract the oxidative stress induced by MET and chytrids. Further studies should examine the role of additional antioxidant mechanisms, such as catalase, glutathione peroxidase, and carotenoids in cyanobacterial defence mechanisms.

In contrast to MET, CB leachate exposure decreased total and intracellular MCs and TBARS levels and increased SOD activity only in uninfected cultures. These results suggest that in the absence of the parasite, CB leachate induces oxidative stress, which is countered by enzymatic and non-enzymatic antioxidant mechanisms such as MCs and SOD. However, under co-exposure to CB leachate and chytrids, these biomarkers remained unaffected suggesting a parasite-mediated mitigation of chemical stress. Cyanobacteria may benefit from parasitic infections [40], as parasites may reduce the impact of pollutants by bioaccumulating them [80,81] or by enhancing host resistance to pollutants through metabolic alterations [82,83].

The IBRv2 was higher in cyanobacteria co-exposed to MET and chytrids, suggesting a synergistic effect. Similar effects have been observed in other host-parasite systems exposed to other pollutants [84–87]. Our results support previous hypotheses that MET may impact chytrids indirectly by altering host nutrient composition without inhibiting host growth. These alterations can impair chytrid fitness over time, as previously observed [39], thereby disrupting key parasite traits, such as nutrient uptake, membrane formation, and energy storage [12,88–90].

Conversely, CB leachate had the opposite effect. IBRv2 was higher in uninfected cyanobacteria but remained unchanged in those co-exposed to CB leachate and chytrids, suggesting an antagonistic effect, where parasites enhance host tolerance to chemical stress. Similar indirect benefits of parasites in pollutant-exposed hosts have been reported for other systems [80–83,91]. Given the complex composition of CB leachate, we cannot pinpoint the observed effects to any specific compound but rather to the mixture as a whole.

Overall, our results support both hypotheses: 1) MET and chytrids act synergistically to increase cyanobacterial stress, and 2) CB leachate exerts a stronger negative effect on uninfected cyanobacteria, with chytrids mitigating this stress.

## 5. Conclusions

Our study highlights the complexity of cyanobacterial responses to simultaneous abiotic (chemical) and biotic (chytrid parasites) stressors, providing mechanistic insights into cyanobacterial-chytrid interactions under pollutant stress. Chytrid parasites, and possibly other phytoplanktonic parasites, can modulate host stress responses, ultimately influencing how pollutants impact aquatic ecosystems. Thus, our results emphasize the need to consider biotic interactions, particularly host-parasite dynamics, in ecotoxicological assessments of cyanobacteria and other phytoplankton to better understand the effects of pollution on ecosystem health.

## Acknowledgments

This work was supported by the Deutsche Forschungsgemeinschaft (DFG) grant MA 9934/1-1. We thank Claudia Schmalsch for her technical support with metolachlor, nicotine and cotinine analyses. The authors declare no competing financial interest.

## Data availability statement

The data that supports the findings of this study and the code used to analyze it are available in the supplementary material of this article (Supplementary material 2).

## Authors’ contribution

**Erika B. Martínez-Ruiz**: Conceptualization, Methodology, Investigation, Formal Analysis, Resources, Writing: Original draft, Review & Editing, Visualization, Supervision, Project administration, Funding acquisition. **Jutta Fastner**: Methodology (Microcystin quantification), Resources, Writing: Review & Editing**. Stephanie Spahr**: Methodology (Chemical analyses), Resources, Writing: Review & Editing. **Justyna Wolinska**: Resources, Writing: Review & Editing.

