## Supplementary material for "Synergistic and antagonistic effects of chemical pollutants and parasitic fungi on cyanobacterial metabolism"

Table S1. Cyanobacterial and chytrid biovolume in uninfected and infected cyanobacterial cultures ($n$ = 5).

| Condition | Treatment | Cyanobacterial biovolume ($mm^3 L^{-1}$) | Chytrid biovolume ($mm^3 L^{-1}$) | Total biovolume ($mm^3 L^{-1}$) | % Cyanob. | % Chytrid |
|---|---|---|---|---|---|---|
| Uninfected | Control | 68.53 | 0.00 | 68.53 | 100.0 | 0.0 |
| Uninfected | MET | 80.94 | 0.00 | 80.94 | 100.0 | 0.0 |
| Uninfected | CB | 90.29 | 0.00 | 90.29 | 100.0 | 0.0 |
| Infected | Control | 117.14 | 0.82 | 117.96 | 99.3 | 0.7 |
| Infected | MET | 164.70 | 1.27 | 165.97 | 99.2 | 0.8 |
| Infected | CB | 120.77 | 0.78 | 121.56 | 99.4 | 0.6 |

Table S2. Linear effect models with interaction predicting metolachlor, nicotine and cotinine concentrations, as well as IBRv2. Estimates significant at the $p < 0.05$ level are bold.

| Effect | Estimate (± s.e.) | *t*-value | *F* | *p-value* |
|---|---|---|---|---|
| Metolachlor | df = 16; adj. $R^2$ = 0.32 | | | |
| Intercept | 9.5 ± 2.18 | 4.35 | | |
| Day × Condition | | | 0.13 | 0.72 |
| Day | | | 3.03 | 0.1 |
| **Condition** | | | **9.4** | **0.007** |
| Nicotine | df = 16; adj. $R^2$ = 0.42 | | | |
| Intercept | 13.6 ± 2.02 | 6.75 | | |
| Day × Condition | | | 0.01 | 0.92 |
| **Day** | | | **16.74** | **0.0008** |
| Condition | | | 1.05 | 0.32 |
| Cotinine | df = 12; adj. $R^2$ = 0.81 | | | |
| Intercept | 5.1 ± 1.15 | 4.42 | | |
| Day × Condition | | | 3.32 | 0.09 |
| **Day** | | | **64.86** | **< 0.0001** |
| **Condition** | | | **5.96** | **0.026** |
| IBRv2 | df = 24; adj. $R^2$ = 0.45 | | | |
| Intercept | 2.7 ± 0.31 | 8.67 | | |
| **Treatment × Condition** | | | **7.32** | **0.003** |

$^+$: F test performed on nested models. In models where the interaction was significant, we did not test the significance of individual fixed effects.

Table S3. Linear effect models predicting cyanobacteria biovolume.

| Effect | Estimate (± s.e.) | Variance (± s.d.) | Degrees of freedom | $t$-value |
|---|---|---|---|---|
| **Cyanobacteria biovolume of uninfected cultures (cond. $R^2$ = 0.309; marg. $R^2$ = 0.073)[a]** | | | | |
| Intercept | 8.03 ± 0.49 | | 12 | 16.43 |
| MET | 0.81 ± 0.69 | | 12 | 1.164 |
| CB | 1.29 ± 0.69 | | 12 | 1.867 |
| Sample | | 0.92 ± 0.96 | | |
| Residual | | 2.71 ± 1.65 | | |
| **Cyanobacteria biovolume of infected cultures (cond. $R^2$ = 0.330; marg. $R^2$ = 0.159)[a]** | | | | |
| Intercept | 10.60 ± 0.51 | | 12 | 20.65 |
| MET | 2.08 ± 0.73 | | 12 | 2.86 |
| CB | 0.21 ± 0.73 | | 12 | 0.28 |
| Sample | | 0.95 ± 0.97 | | |
| Residual | | 3.70 ± 1.93 | | |

[a]Cond. $R^2$: conditional $R^2$ describes the proportion of total variance explained by the fixed effects and the random effects together; marg. $R^2$: marginal $R^2$ describes the proportion of total variance explained exclusively by the fixed effects. Root square transform data was used for the models.

Table S4. Linear effect models predicting infection prevalence, infection intensity and biomarker responses.

| | **Infection prevalence (df = 12; adj. $R^2$ = 0.65)** | | **Infection intensity (df = 12; adj. $R^2$ = 0.39)** | | **Proteins uninfected (df = 12; adj. $R^2$ = 0.40)** | | **Proteins infected (df = 12; adj. $R^2$ = 0.42)** | |
|---|---|---|---|---|---|---|---|---|
| Effect | Estimate (± s.e.) | *t*-value | Estimate (± s.e.) | *t*-value | Estimate (± s.e.) | *t*-value | Estimate (± s.e.) | *t*-value |
| Intercept | 9.7 ± 1.19 | 8.17 | 7.3 ± 1.54 | 4.74 | 2.80 ± 0.86 | 32.61 | 9.2 ± 1.53 | 6.03 |
| MET | 1.6 ± 1.68 | 0.95 | 4.6 ± 2.18 | 2.11 | -0.26 ± 0.12 | -2.12 | -5.4 ± 2.16 | -2.50 |
| CB | -6.7 ± 1.68 | -3.99 | -2.5 ± 2.18 | -1.15 | -0.4 ± 0.12 | -3.30 | 1.8 ± 2.16 | 0.83 |

Table S4. (continued)

| | **Lipids uninfected (df = 12; adj. $R^2$ = 0.47)** | | **Lipid infected (df = 12; adj. $R^2$ = 0.21)** | | **TBARS uninfected (df = 12; adj. $R^2$ = 0.41)** | | **TBARS infected (df = 12; adj. $R^2$ = 0.42)** | |
|---|---|---|---|---|---|---|---|---|
| Effect | Estimate (± s.e.) | *t*-value | Estimate (± s.e.) | *t*-value | Estimate (± s.e.) | *t*-value | Estimate (± s.e.) | *t*-value |
| Intercept | 11.8 ± 1.46 | 8.09 | 11.0 ± 1.78 | 6.18 | 12.2 ± 1.54 | 7.91 | 11.4 ± 1.53 | 7.45 |
| MET | -3.6 ± 2.06 | -1.75 | -6.0 ± 2.52 | -2.38 | -5.6 ± 2.18 | -2.57 | -7.4 ± 2.16 | -3.42 |
| CB | -7.8 ± 2.06 | -3.79 | -3.0 ± 2.52 | -1.19 | -7.0 ± 2.18 | -3.21 | -2.8 ± 2.16 | -1.29 |

Table S4. (continued)

| | **SOD uninfected (df = 12; adj. $R^2$ = 0.48)** | | **SOD infected (df = 12; adj. $R^2$ = -0.06)** | | **GST uninfected (df = 12; adj. $R^2$ = -0.06)** | | **GST infected (df = 12; adj. $R^2$ = -0.12)** | |
|---|---|---|---|---|---|---|---|---|
| Effect | Estimate (± s.e.) | *t*-value | Estimate (± s.e.) | *t*-value | Estimate (± s.e.) | *t*-value | Estimate (± s.e.) | *t*-value |
| Intercept | 4.8 ± 1.44 | 3.33 | 8.6 ± 2.06 | 4.17 | 6.2 ± 2.06 | 3.01 | 8.6 ± 2.12 | 4.06 |
| MET | 2.0 ± 2.04 | 0.98 | -2.4 ± 2.91 | -0.82 | 3.0 ± 2.91 | 1.03 | 6.08 ± 3.00 | 0 |
| CB | 7.6 ± 2.04 | 3.73 | 0.60 ± 2.91 | 0.21 | 2.4 ± 2.91 | 0.82 | -1.8 ± 3.00 | -0.60 |

Table S5. Percentage of microcystins in infected and uninfected cultures. Values represent the mean ± sd, (*n* = 5).

| Status | Treatment | DMC-RR (%) | DMC-LR (%) | MC X (%) |
|---|---|---|---|---|
| Uninfected | Control | 90.7 ± 0.73 | 8.6 ± 0.73 | 0.7 ± 0.02 |
| Uninfected | MET | 93.1 ± 1.43 | 6.4 ± 1.36 | 0.5 ± 0.07 |
| Uninfected | CB | 91.2 ± 0.60 | 8.2 ± 0.61 | 0.7 ± 0.02 |
| Infected | Control | 94.6 ± 0.30 | 4.9 ± 0.32 | 0.5 ± 0.04 |
| Infected | MET | 91.2 ± 1.06 | 8.3 ± 1.08 | 0.6 ± 0.02 |
| Infected | CB | 95.0 ± 0.25 | 4.6 ± 0.24 | 0.4 ± 0.04 |

Table S6. Linear effect models predicting MC concentrations

.

| Effect | Uninfected | | Infected | |
|---|---|---|---|---|
| | Estimate (± s.e.) | *t*-value | Estimate (± s.e.) | *t*-value |
| **DMC-RR total** | **df = 12; adj. $R^2$ = 0.34** | | **df = 12; adj. $R^2$ = 0.55** | |
| Intercept | 11 ± 1.63 | 6.75 | 9.2 ± 1.35 | 6.82 |
| MET | -2.2 ± 2.31 | -0.95 | -5.8 ± 1.91 | -3.04 |
| CB | -6.8 ± 2.31 | -2.95 | 2.20 ±1.91 | 0.27 |
| **DMC-RR intracellular** | **df = 12; adj. $R^2$ = 0.34** | | **df = 12; adj. $R^2$ = 0.55** | |
| Intercept | 11.0 ± 1.63 | 6.75 | 9.2 ± 1.35 | 6.82 |
| MET | -2.2 ± 2.31 | -0.95 | -5.8 ± 1.91 | -3.04 |
| CB | -6.8 ± 2.31 | -2.95 | 2.2 ± 1.91 | 1.15 |
| **DMC-RR extracelullar** | **df = 12; adj. $R^2$ = 0.24** | | **df = 12; adj. $R^2$ = 0.44** | |
| Intercept | 11.2 ± 1.75 | 6.4 | 11.8 ± 1.50 | 7.85 |
| MET | -3.4 ± 2.47 | -1.37 | -3.8 ± 2.13 | -1.79 |
| CB | -6.2 ± 2.47 | -2.51 | -7.6 ± 2.13 | -3.58 |
| **DMC-LR total** | **df = 12; adj. $R^2$ = 0.42** | | **df = 12; adj. $R^2$ = 0.50** | |
| Intercept | 11.0 ± 1.53 | 7.21 | 0.34 ± 0.03 | 12.19 |
| MET | -1.8 ± 2.16 | -0.83 | -0.16 ± 0.04 | -4.02 |
| CB | -7.2 ± 2.16 | -3.34 | -0.08 ± 0.04 | -2.01 |
| **DMC-LR intracellular** | **df = 12; adj. $R^2$ = 0.42** | | **df = 12; adj. $R^2$ = 0.48** | |
| Intercept | 11.0 ± 1.53 | 7.21 | 0.34 ± 0.03 | 11.94 |
| MET | -1.8 ± 2.16 | -0.83 | -0.16 ± 0.04 | -3.89 |
| CB | -7.2 ± 2.16 | -3.34 | -0.08 ± 0.04 | -1.91 |
| **DMC-LR extracelullar** | **df = 12; adj. $R^2$ = 0.065** | | **df = 12; adj. $R^2$ = 0.41** | |
| Intercept | 10.6 ± 1.93 | 5.48 | 11.0 ± 1.08 | 10.22 |
| MET | -3.2 ± 2.74 | -1.17 | -4.5 ± 1.52 | -2.96 |
| CB | -4.6 ± 2.74 | -1.68 | -4.5 ± 1.52 | -2.96 |

| | | | | |
|---|---|---|---|---|
| **MC X total** | **df = 12; adj. $R^2$ = 0.47** | | **df = 12; adj. $R^2$ = 0.22** | |
| Intercept | 11.0 ± 1.46 | 7.52 | 10.4 ± 5.87 | 5.87 |
| MET | -1.6 ± 2.07 | -0.77 | -5.8 ± 2.51 | -2.31 |
| CB | -7.4 ± 2.07 | -3.58 | -1.4 ± 2.51 | -0.56 |
| **MC X intracelullar** | **df = 12; adj. $R^2$ = 0.47** | | **df = 12; adj. $R^2$ = 0.22** | |
| Intercept | 11.0 ± 1.46 | 7.52 | 10.4 ± 1.77 | 5.87 |
| MET | -1.6 ± 2.07 | -0.77 | -5.8 ± 2.51 | -2.31 |
| CB | -7.4 ± 2.07 | -3.58 | -1.4 ± 2.51 | -0.56 |
| **MC X extracelullar** | **df = 12; adj. $R^2$ = -0.12** | | all data points were below the detection limit | |
| Intercept | 7.6 ± 1.65 | 4.62 | | |
| MET | 1.4 ± 2.33 | 0.60 | | |
| CB | -2.0 ± 2.33 | -0.09 | | |

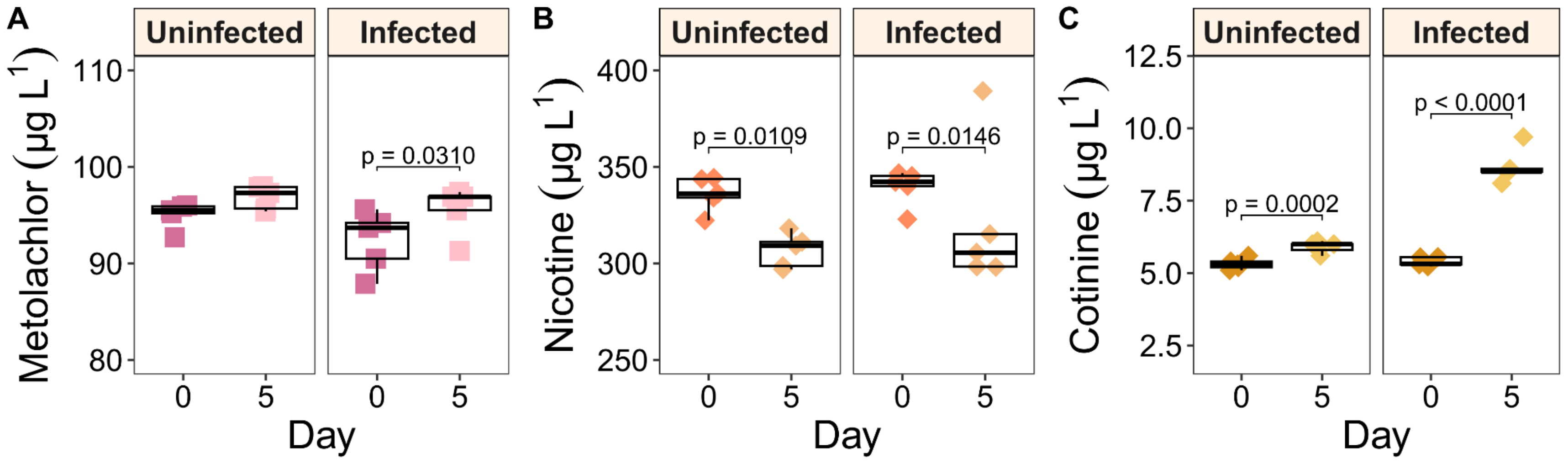


Figure S1. Metolachlor, nicotine and cotinine concentrations in cyanobacteria exposed to chemical pollutants with and without chytrid parasites. Boxes indicate upper and lower quartiles, the dark middle line indicates the median, whiskers indicate 1.5 times the interquartile range and black points represent outliers ($n$ = 5).

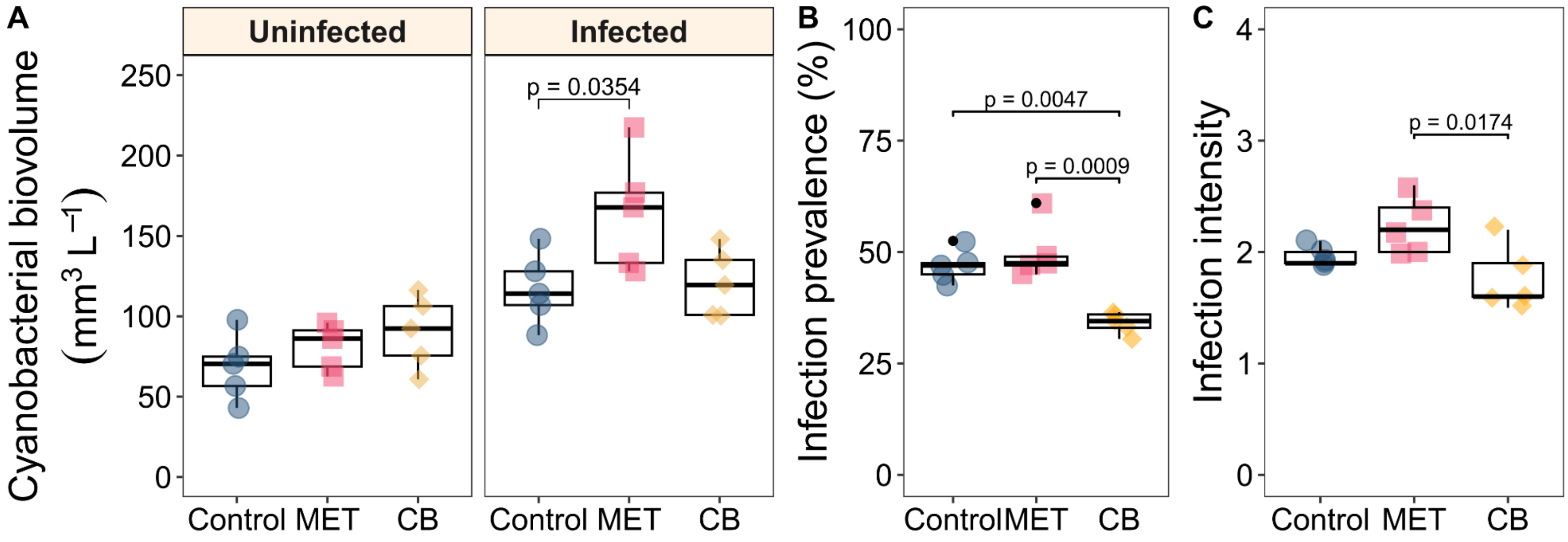


Figure S2. Cyanobacterial biovolume and parasite fitness proxies. A) Cyanobacterial biovolume, B) infection prevalence (percentage of infected cyanobacterial filaments), and C) infection intensity (mean infections number per infected filament). Boxes indicate the upper and lower quartiles, the dark middle line indicates the median, whiskers indicate 1.5 times the interquartile range and black points represent outliers ($n$ = 5). MET: metolachlor; CB: cigarette butt leachate.

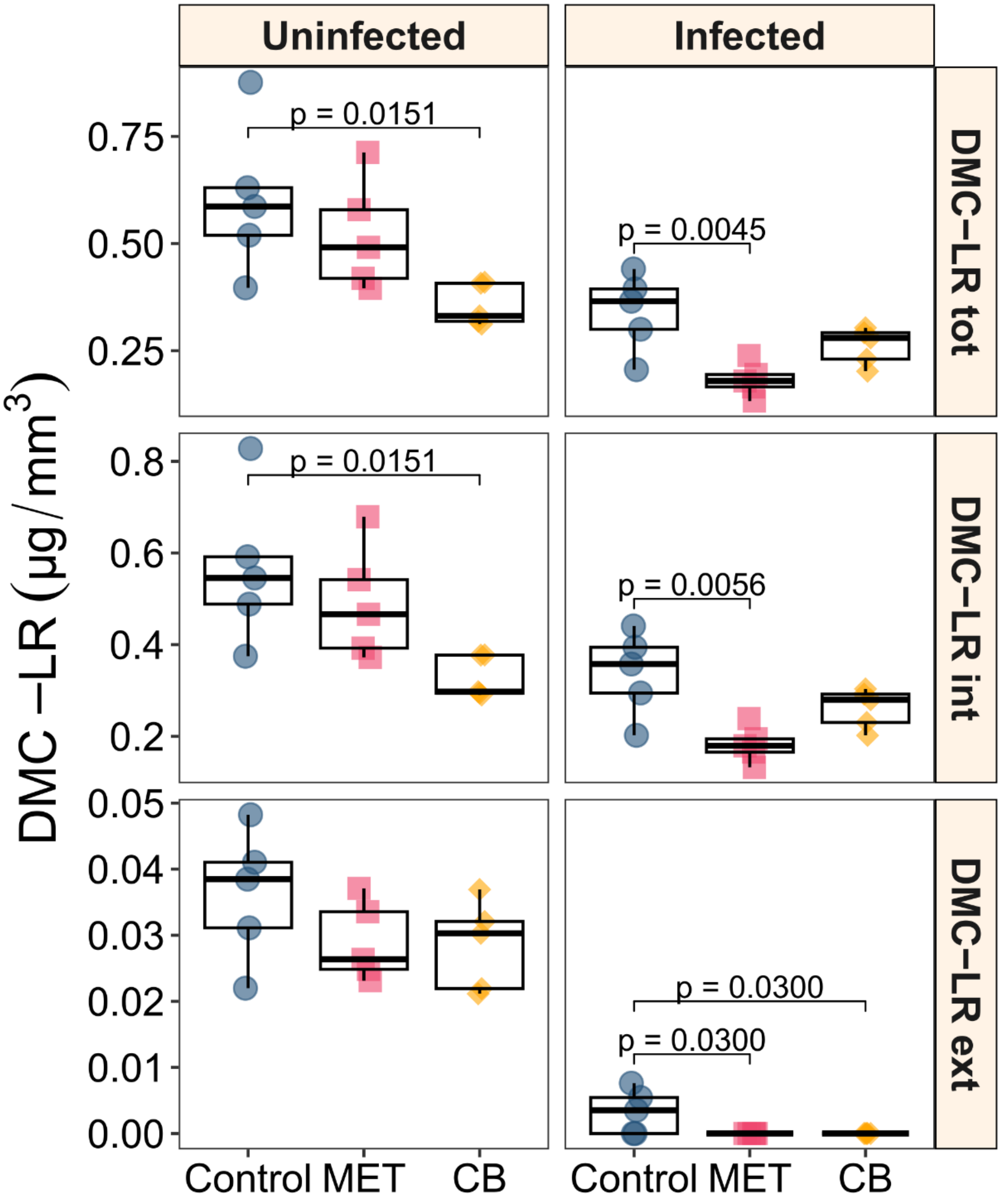


Figure S3. Total, intracellular and extracellular DMC-LR concentrations in cyanobacteria exposed to chemical pollutants with and without chytrid parasites. DMC-LR concentrations were adjusted to the cyanobacterial biovolume of each replicate per treatment and condition. Boxes indicate the upper and lower quartiles, the dark middle line indicates the median, whiskers indicate 1.5 times the interquartile range and black points represent outliers ($n$ = 5). MET: metolachlor; CB: cigarette butt leachate; DMC-LR tot: MC-LR total; DMC-LR int: MC-LR intracellular and DMC-LR ext: MC-LR extracellular.

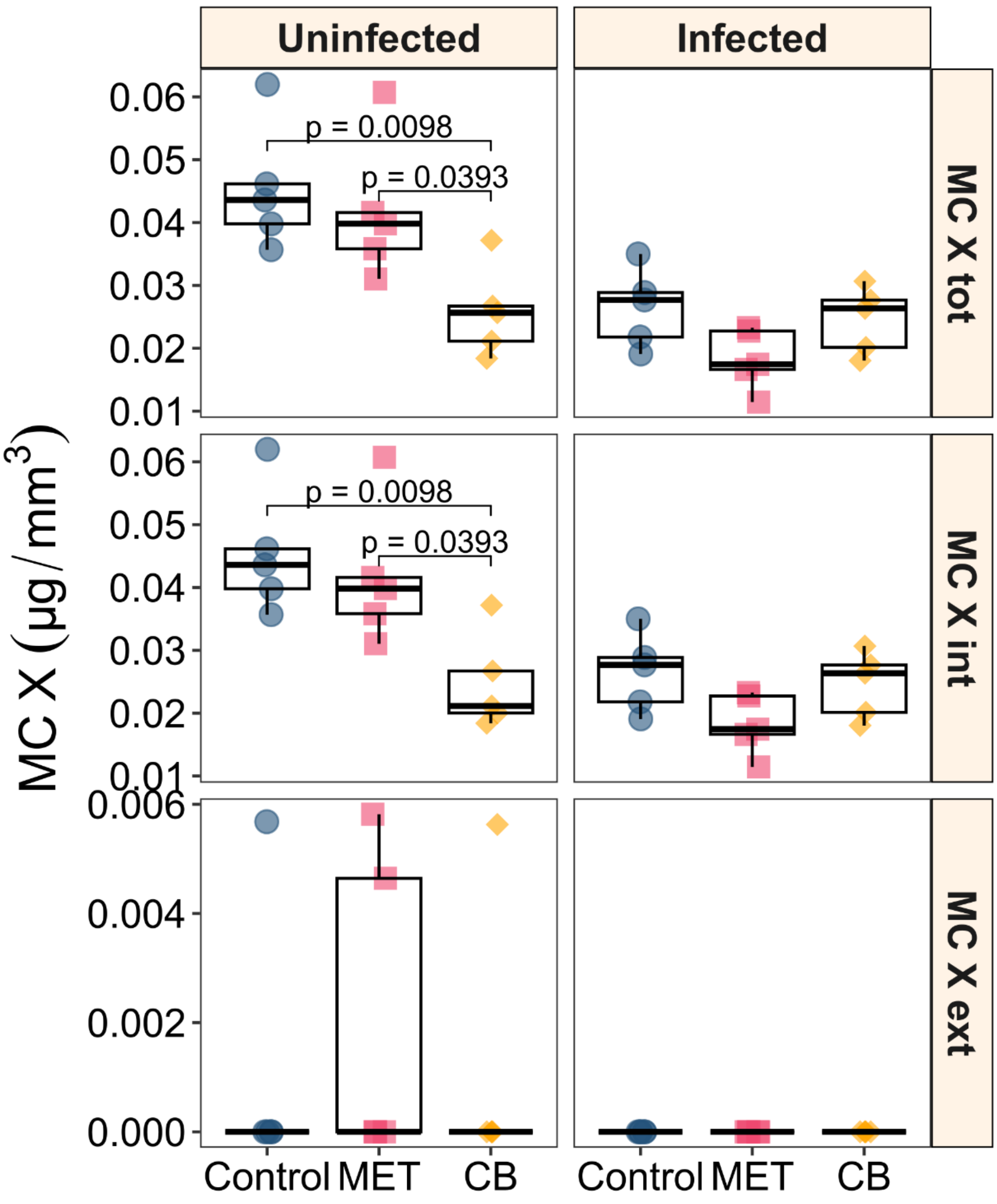


Figure S4. Total, intracellular and extracellular MC X concentrations in cyanobacteria exposed to chemical pollutants with and without chytrid parasites. MC X concentrations were adjusted to the cyanobacterial biovolume of each replicate per treatment and condition. Boxes indicate upper and lower quartiles, the dark middle line indicates the median, whiskers indicate 1.5 times the interquartile range and black points represent outliers ($n$ = 5). MET: metolachlor; CB: cigarette butt leachate; MC X tot: MC X total; MC X int: MC X intracellular; MC X ext: MC X extracellular.

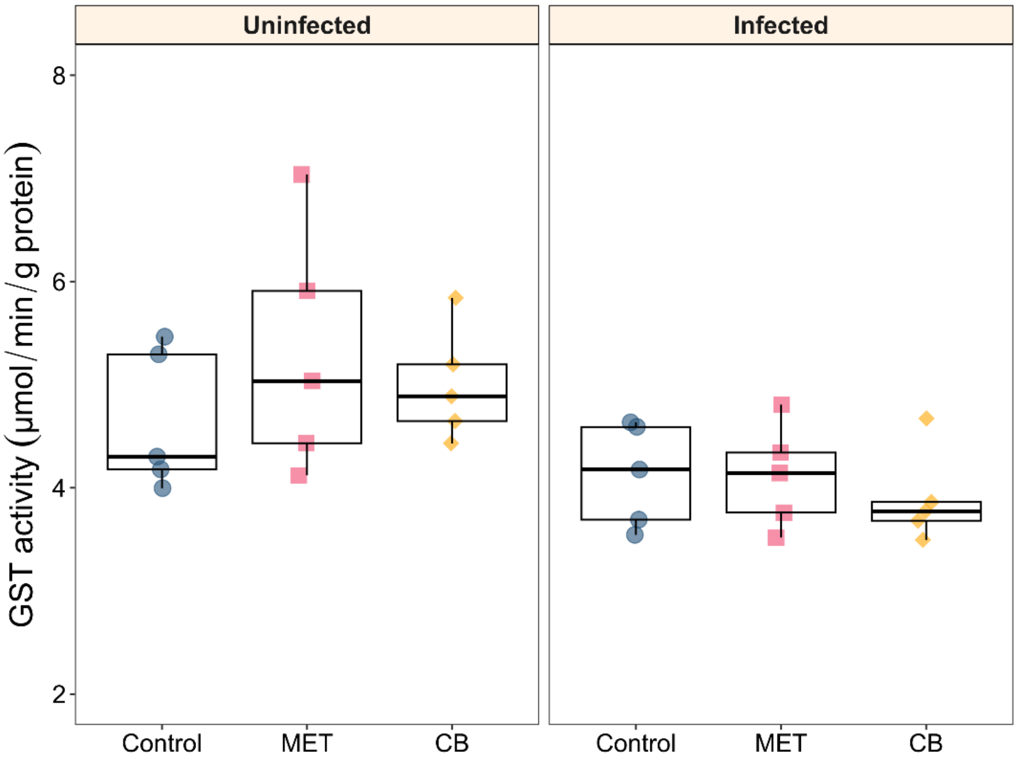


Figure S5. Glutathione *S*-transferase activity in cyanobacteria exposed to chemical pollutants with and without chytrid parasites. GST activity was adjusted to the protein content of each replicate per treatment and condition. Boxes indicate the upper and lower quartiles, the dark middle line indicates the median, whiskers indicate 1.5 times the interquartile range and black points represent outliers ($n$ = 5). MET: metolachlor; CB: cigarette butt leachate; GST: glutathione *S*-transferase.
